# Transcriptome Sequencing Reveals Sex Differences in Human Meniscal Cell Response to Estrogen Based on Dosing Kinetics

**DOI:** 10.1101/2020.04.27.064451

**Authors:** Kelsey E. Knewtson, Jesus G. Gonzalez Flores, Donna M. Pacicca, Jennifer L. Robinson

**Author notes:** **Corresponding Author** Jennifer L. Robinson, PhD, 1530 West 15^th^ Street Room 4132, Lawrence, KS 66045. **Author Contributions** K.K. analyzed the data and wrote the manuscript. J.G.F. designed and performed the experiment and reviewed the manuscript. D.P. provided critical interpretation of the data and revised the manuscript. J.R. designed the experiment, supervised the study, and wrote the manuscript. All authors have read and approved of the submitted manuscript.

## Abstract

Osteoarthritis is a disease marked by progressive and irreversible hyaline cartilage and fibrocartilage breakdown that affects the lives of millions of patients worldwide. Female sex and menopause are both risk factors for knee osteoarthritis, indicating that estrogen could play a role in this disease. In this study, RNA sequencing was used to determine the effects of estrogen treatment on human meniscal cells. Differences in the number and type of differentially expressed genes were seen based on donor sex, estrogen dose, and dosing kinetics. Significantly more differentially expressed genes were seen from male meniscal cells in response to all dosing conditions compared to female cells. Importantly, more genes were differentially expressed in cells treated with continuous dosing of estrogen, which has been shown to stimulate genomic estrogen signaling, as compared to pulsed dosing. Additionally, functional enrichment analysis revealed that many genes of the extracellular matrix, which is important for joint health and injury repair, were differentially expressed. Overall, this initial study lays the groundwork for future avenues to pursue the effect of estrogen delivery on regenerative pathways. This critical analysis will then inform the design and implementation of estrogen replacement therapies to promote meniscal health and reduce the onset of osteoarthritis.

## INTRODUCTION

An estimated 250 million people have knee osteoarthritis (OA) worldwide,^1^ with more than 25 million cases in the United States alone.^2^ Patients experience progressive and irreversible cartilage breakdown that leads to pain, immobility, and eventual joint failure. Meniscal tears are the most prevalent intra-articular knee injury and pose significant risk in the development of OA.^3^ Knee OA is more common and severe in women than in men, especially post-menopause.^4^ Also, tears of the meniscal body are more common in women than in men.^5^ Further, recent epidemiological studies have found that meniscal injury rates are higher in female high school athletes than in males competing in comparable sports,^6^ and that females require repeat meniscal repair surgeries 2.2x more frequently.^7^ These data suggest that sex hormone levels, including estrogen, may play a role in meniscal injury and repair.

Estrogen is required for joint health and is used as a therapeutic for joint degenerative conditions. Several studies have found a protective effect of estrogen replacement therapy (ERT) against the development and progression of knee OA in post-menopausal women; however, the effects were modest and did not reach significance.^8–10^ Treatment with estrogen or selective estrogen receptor modulators has been shown to improve OA symptoms in humans in some studies, but the results are mixed and inconclusive.^11^ While clinical data suggests estrogen protects the knee joint and reduces OA, there is a dearth of *in vitro* and preclinical studies investigating the role of estrogen on meniscal health and repair. A better understanding of the effects of estrogen treatment on the knee is needed to develop therapies to improve joint health in different populations.

The physiological effects of estrogen vary depending on concentration and dosing kinetics. High and low concentrations of estrogen are known to have opposite effects on inflammation,^12^ gene expression,^13^ and stem cell proliferation.^14^ Specific 17β-estradiol (E2) effects are dose-dependent and differ between tissues.^15^ Additionally, several studies have found that the effect of estrogen varies based on dosing kinetics both *in vitro*^*16; 17*^ and *in vivo*.^18^ Typically, estrogen exerts its effects on cells by activating the nuclear estrogen receptors alpha and beta, which then regulate transcription of many genes. However, it is known that estrogen can also have quicker, non-genomic effects on cells.^19^ There is evidence that controlling dosing kinetics can control the type of response elicited.^16^ Conversely, others have found that if total hormone exposure is kept constant, dosing kinetics do not alter the effects of estrogen.^20^

The intricacies of the effects of estrogen are significant given the different levels of estrogen seen in patients of different sexes and stages of life and the different doses and delivery routes used for ERT. Serum estradiol levels in healthy, menstruating women range from 40-400 pg/mL (150-1,500 pM) depending on the phase of the menstrual cycle,^21^ while levels in pregnant women range from 2,000-20,000 pg/mL (7,300-73,000 pM) depending on trimester.^22^ Post-menopausal women typically have serum estradiol levels below 15 pg/mL (60 pM).^21^ Healthy men under age 55 average serum estradiol levels less than 40 pg/mL (150 pM),^21; 23^ and these levels decrease as men age and their testosterone levels decrease.^23^ Average serum levels of estradiol achieved by ERT typically range from 20-100 pg/mL.^24^ Less data is available on synovial fluid (SF) estradiol levels, which is the most pertinent data for tissues such as the meniscus. The data available for SF is mostly limited to older patient populations with OA or rheumatoid arthritis (RA). These studies have found estradiol SF levels ranging from 15-54 pg/mL (60-200 pM) in male and female patients, with higher levels typically seen in RA patients than in OA patients.^25; 26^ Though these levels cannot be applied to other populations, these studies did find strong, positive linear correlations between serum and SF estradiol levels that could be applicable to other groups. In terms of ERTs, each treatment type (e.g. formulation and deployment form) is associated with a different hormone exposure level and kinetic profile. For example, though a 50 μg/day transdermal reservoir patch, 2 mg oral tablet, and 300 μg nasal dose of E2 all provide similar 24 h systemic exposures (1080-1400 pg*h/ml), they each have a unique kinetic profile ranging from continuous over days with a transdermal patch to pulsed over hours with a nasal delivery route.^27^ Further studies are needed to understand the effects of estrogen concentration and dosing kinetics on tissues of interest from different patient populations.

In this study, we aimed to both determine sex differences and the role of concentration and dosing kinetics on human knee meniscal cell gene expression changes in response to estrogen treatment. Cells were isolated from the menisci of two age-matched donors, one of each sex, and treated with two concentrations of 17β-estradiol (E2) under pulsed and continuous conditions. Messenger RNA sequencing (RNA-Seq) was then used to determine differentially expressed genes as a function of the E2 treatment groups. Results from this study will inform future work on developing a mechanistic understanding of estrogen delivery conditions on cell and tissue response to aid in design and development of treatments for knee injury based on patient sex.

## METHODS

### Cells and Treatment

The experimental design and workflow are illustrated in Figure 1. Human meniscal tissue was received from the National Disease Research Interchange (NDRI). Cells were harvested from the medial meniscus of a human cadaveric male (Black) and a female (Caucasian) (both 47-year old). Briefly, whole menisci were cut into small pieces (1-2 mm) and digested in DMEM (11996-065, Gibco, Carlsbad, CA) containing 2% penicillin/streptomycin (15140-122, Gibco, Carlsbad, CA) and 3 mg/mL collagenase type I (17100-017, Gibco, Carlsbad, CA) at 37 °C and 60 rpm for 16 h. Solutions were filtered through Swinnex Filter Holders (25 mm, SX0002500, Sigma-Aldrich, St. Louis, MO) loaded with Spectra Mesh Nylon Filters (30 μM, 888-13681, Spectrum Chemical Manufacturing Corporation, New Brunswick, NJ) to remove undigested tissue. Cells were then cultured in T75 tissue culture treated flasks in DMEM supplemented with 10% Premium FBS (511150, Atlanta Biologicals, Minneapolis, MN) and 1% penicillin/streptomycin. After reaching confluency, cells were passaged using trypsin (T4174, Sigma, St. Louis, MO, diluted to 1x before use) and seeded into T225 tissue culture treated flasks. When passage 1 cells reached 70-80% confluency, cells were washed with PBS (20012-207, Gibco, Carlsbad, CA) and cultured in phenol red-free DMEM (pfDMEM, 31053-028, Gibco, Carlsbad, CA) with 15% charcoal/dextran treated FBS (S11650, Atlanta Biologicals, Minneapolis, MN) for 3 days to starve them of endogenous hormones. Cells were then passaged and seeded at 10,000 cells/cm^2^ in 24 well plates in pfDMEM and allowed to adhere overnight. The next day, cells were treated in triplicate with 17β-estradiol (E2, E2758, Sigma, St. Louis, MO) in pfDMEM supplemented with 15% charcoal/dextran treated FBS and 1% penicillin/streptomycin under pulsed or continuous dosing. For pulsed dosing, cells were exposed to E2 (0, 24, or 240 μM) for 1 h, after which the E2 media was replaced with E2-free media for 23 h. This pattern was repeated for a total of 72 h. For continuous dosing, cells were exposed to E2 (0, 1, or 10 μM) for the entire treatment period of 72 h. The different concentrations used for pulsed compared to continuous dosing were used so that the total 24 h exposure was identical between groups, as reported previously.^16; 17; 20^ The concentrations used were chosen to be above normal physiological levels in order to elicit a significant response. Further, 1 μM is predominantly used as a high concentration in *in vitro* estrogen dosing studies. Different dosing kinetics were used to mimic those seen with different types of estrogen replacement therapies such as pulsed from nasal doses of E2 and continuous from transdermal patches.

**Figure 1:**
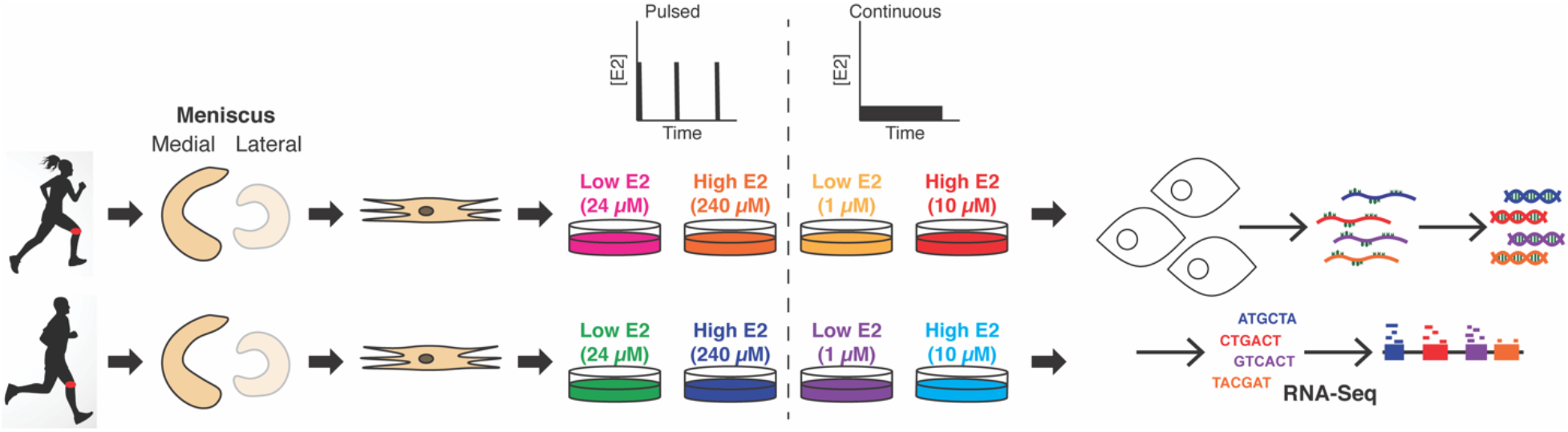
Experimental design. Cells were isolated from the medial meniscus of two 47-year old donors, one of each sex, and treated with two concentrations of E2 under pulsed and continuous conditions. RNA-Seq was then used to analyze the samples. Running silhouettes from freepik.com.

### RNA Extraction, cDNA Library Construction, and Sequencing

After 72 h, cells were collected in TRIzol, and total RNA was purified using the manufacturer’s protocol. DNA was removed using a DNA-*free* DNA Removal Kit (AM1906, Life Technologies, Grand Island, NY). Total RNA samples were quality checked with a Qubit Assay and Agilent TapeStation for quantity and quality, respectively. All RNA preps had an estimated RIN value that ranged from 8.8-10. Sequencing libraries were constructed using the NEBNext Ultra II Directional RNA Library Prep Kit for Illumina (E7760, New England BioLabs, Ipswich, MA). The library preps were started with 92-1000 ng of total RNA. The sequencing library construction process includes mRNA purification with polyA selection beads, fragmentation to inserts of ~200 bp, strand specific cDNA synthesis, end repair, 3’ end adenylation, adapter ligation, and PCR amplification. The constructed sequencing libraries were validated and quantified with Qubit and TapeStation assays. Each library was indexed with a barcode sequence and sequenced in a multiplexed fashion. An Illumina NextSeq 550 system at the University of Kansas’s Genome Sequencing Core was used to generate single-end, 100-base sequence reads from the libraries.

### Data Processing

Base calling was carried out by the instrument Real Time Analysis (RTA) software. The base call files (bcl files) were demultiplexed and converted to compressed FASTQ files by bcl2fastq2. Raw reads were filtered using fastp,^28^ retaining only reads longer than 40 nucleotides. Processed reads were then pseudoaligned to the human reference transcriptome (ensembl.org, release 97, build GRCh38) using kallisto,^29^ employing 100 bootstraps to estimate transcript abundance. tximport^30^ was then used to import kallisto transcript abundance estimates into DESeq2^31^ for differential expression analysis. Within DESeq2, a series of pairwise likelihood ratio tests to identify genes showing differential gene expression, considering genes to be differentially expressed for a given test if they survived a 5% False Discovery Rate criterion, was conducted. GraphPad Prism 8 was used to generate volcano plots by plotting the log_2_FoldChange against the adjusted P value. Venny 2.1^32^ was used to generate Venn diagrams of the differentially expressed genes with adjusted p values (padj) < 0.05. Venn diagrams were then redrawn using Adobe Illustrator to allow for color control. Significant results were further analyzed using the Search Tool for the Retrieval of Interacting Genes/Proteins (STRING)^33^ to detect protein-protein interactions using the confidence view to display connection thickness based upon strength of the supporting data. Functional enrichment analysis was performed within STRING, with databases and categories selected based upon first significance and then ability to highlight gene clusters. The Database for Annotation, Visualization and Integrated Discovery (DAVID)^34^ was used for further functional enrichment analysis with the human genome used as the background. The Gene Ontology Cellular Component database was used within the Functional Annotation Chart function of DAVID. Heat maps were generated using GenePattern.^35^ To generate the list of extracellular matrix genes, the genes identified within this category by DAVID for each treatment group were compiled into a master list of ECM genes. The log_2_FoldChange for these genes were then gathered from each treatment group and used uploaded to the Heat Map Image module of GenePattern. Column and row sizes were set to 128 pixels, the color scheme was globally normalized, and color gradient was not used.

## RESULTS

### Male Meniscal Cells Respond More Robustly to Estrogen Treatment

The number of genes in each treatment group that were differentially expressed (DEGs) as compared to the sex-matched no E2 control was plotted in terms of E2 concentration relative to physiological concentrations (Figure 2). This revealed that male cells responded more strongly to E2 treatment than female cells. In general, more DEGs were seen from groups treated with continuous dosing than for groups treated with pulsed dosing, though the male high dose pulsed group had the most DEGs of all treatment groups. Additionally, within the female treatment groups, there were greater differences in the number of DEGs based on dosing kinetics than on concentration. In males, E2 concentration had a large effect on DEG count for the pulsed groups, an effect that does not trend with the other comparisons in which concentration did not play as significant a role when controlling for kinetics.

**Figure 2:**
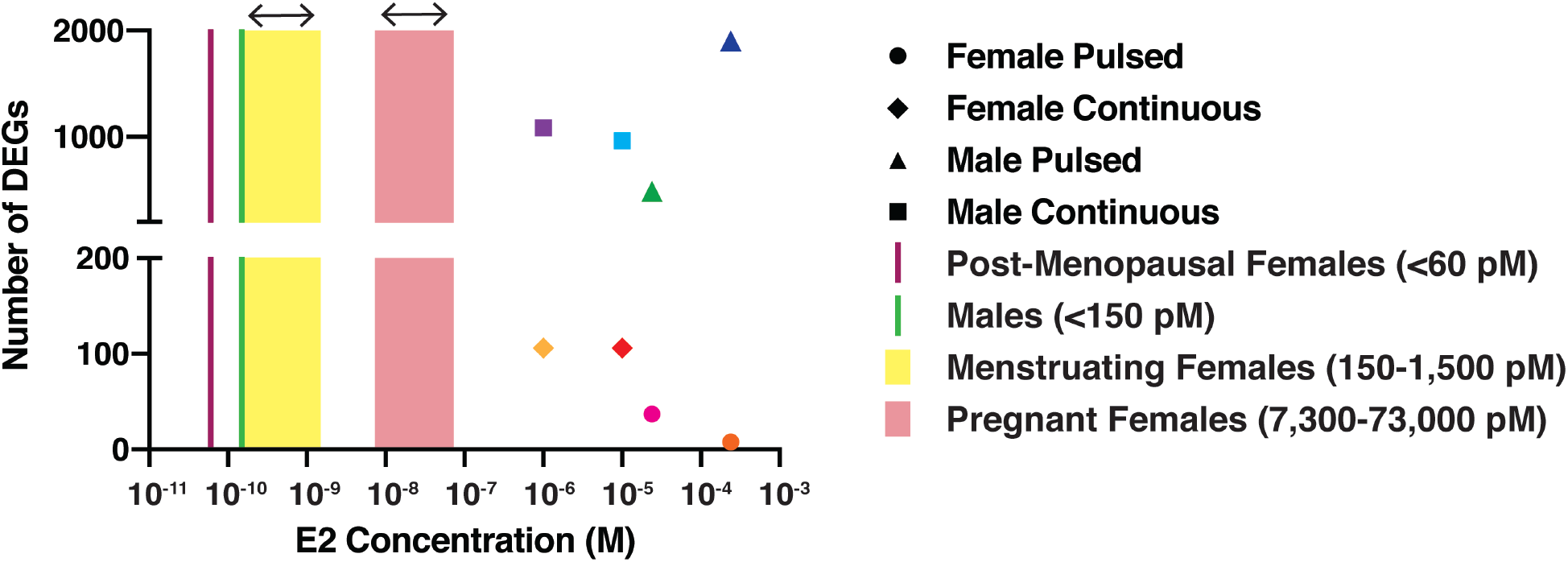
Visualization of the number of differentially expressed genes (DEGs) by treatment group in comparison to physiological serum estrogen concentrations.

Volcano plots were generated to visualize the number of DEGs in each treatment group in terms of fold change and adjusted P value (Figure 3A). These plots show that in most treatment groups, the majority of DEGs had a log_2_FoldChange between −2 and 2, though most groups had at least one gene that fell far outside this range. Examples include POLR2J3 in the female high dose pulsed group (log_2_FoldChange −7.98) and GNL1 in the male high dose pulsed group (log_2_FoldChange 9.94)

**Figure 3:**
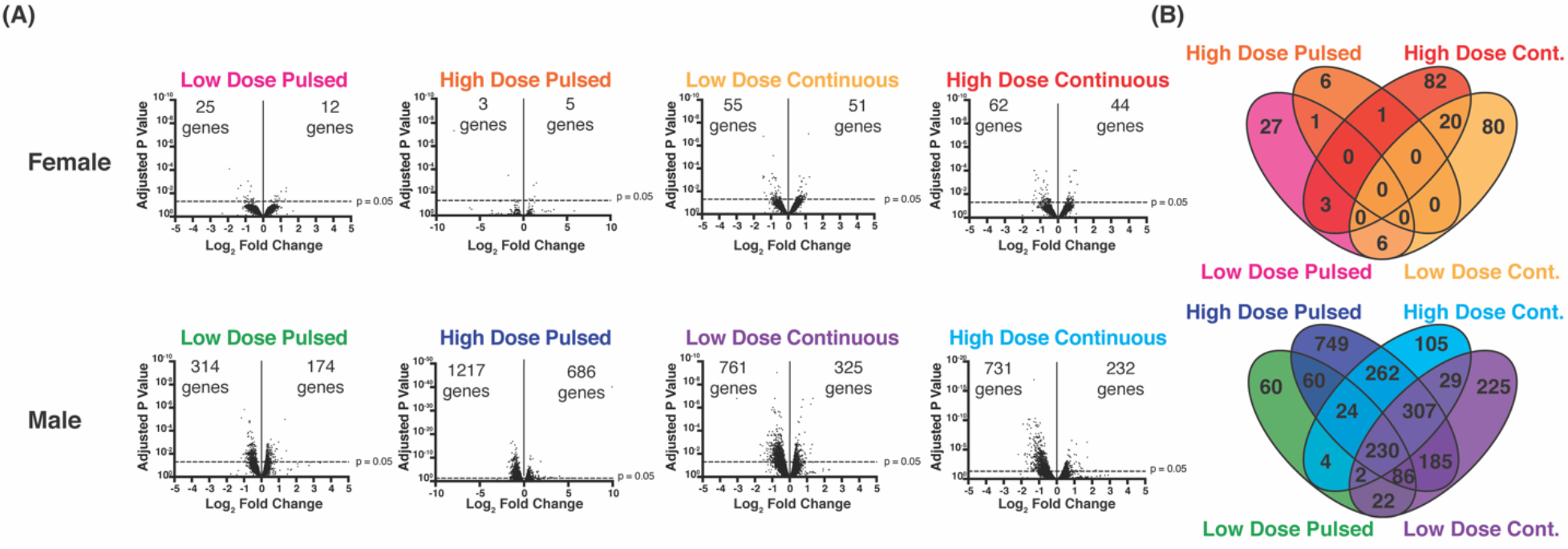
Visualization of differentially expressed genes (DEGs). Volcano plots and Venn diagrams (Venny 2.1) illustrate the number of differentially expressed genes in each treatment group. Only DEGs with padj<0.05 were used in the Venn diagrams.

The Venn diagrams in Figure 3B illustrate more overlap in the DEGs of the male treatment groups compared to the female groups. In fact, 230 genes were common to all 4 groups. Further analysis of these overlapping genes can be seen in Table S1 and Figure S1. This analysis revealed that many groups of genes, including those of the extracellular matrix (ECM), were enriched in this set and that there were many protein-protein interactions among the genes. The highest overlap in DEGs occurred between the male high dose groups, where 823 DEGs were seen in both groups (262 genes that only overlap between the high dose groups and 561 that also overlap with other groups). Comparatively, the highest overlap in female meniscal cell treatment was seen in the continuous treatment groups, where 20 DEGS were seen in both groups.

### Types of DEGs Differ Based on Estrogen Dosing Concentration and Kinetics and Sex

The DEGs from each continuous dosing treatment group were then analyzed using STRING^33^ (Figure 4). Additional constraints based on log_2_FoldChange were used to limit DEGs to a reasonable number for visualization and to highlight those with greater fold changes compared to control. STRING analysis of the other treatment groups and images without log_2_FoldChange constraints can be found in Figures S2-S4. Functional enrichment analysis was also performed within STRING, as indicated by the color coding of certain proteins. This revealed that the DEGs of both female continuous dosing groups were enriched for proteins of the ECM, which is of interest for its importance to joint healing. In the male continuous groups, low dose resulted in DEGs related to metabolic pathways involving nitrogen and high dose resulted in DEGs involved in gene expression and thyroid hormone signaling pathways.

**Figure 4:**
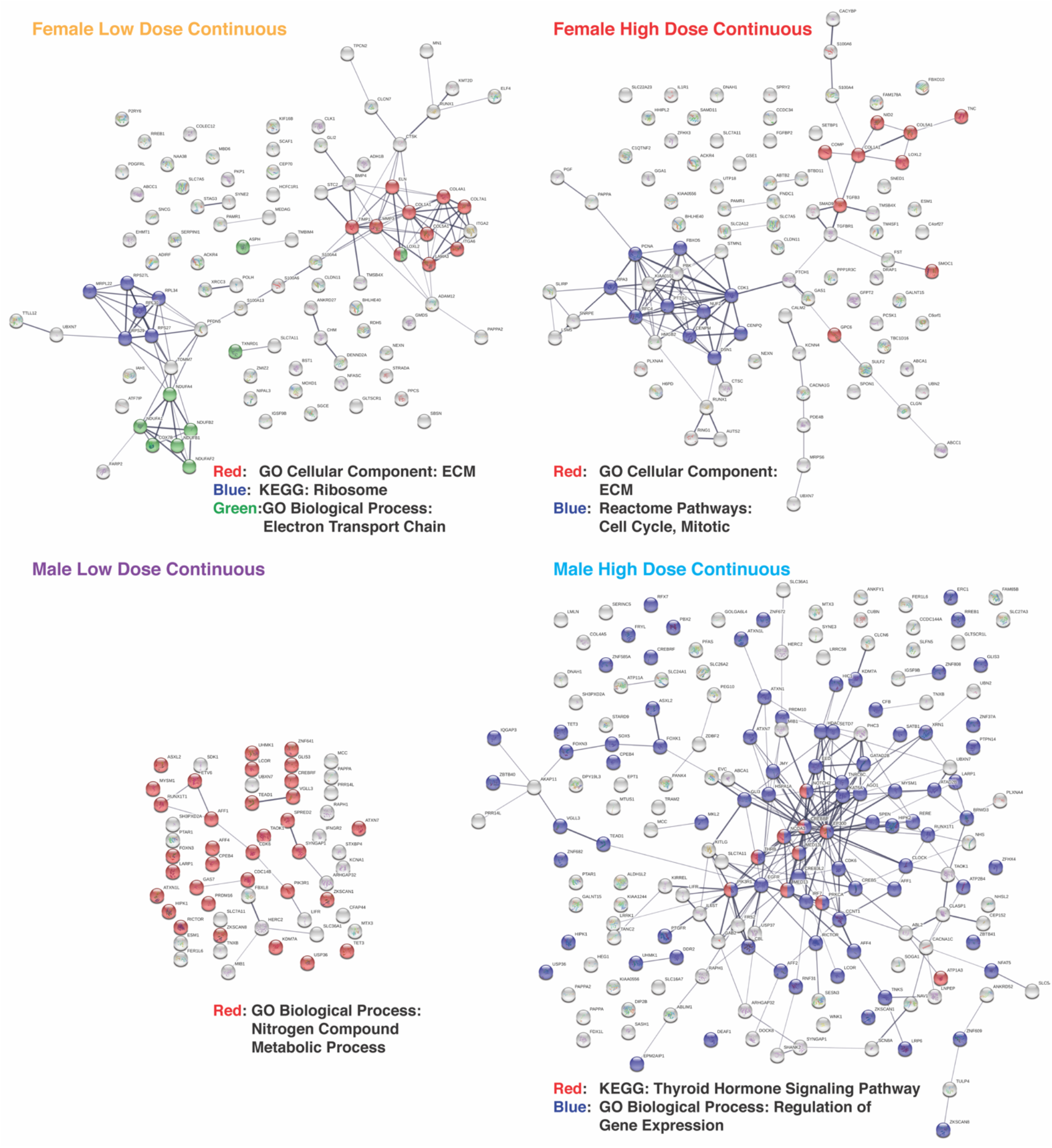
STRING analysis of DEGs from all 4 continuous dosing treatment groups. Images are shown as confidence views, where associations with stronger data support are thicker. Only DEGs with padj<0.05 are shown. Additional constraints based on log_2_FoldChange were used to limit DEGs to a reasonable number for plotting and to highlight those with greater log_2_FoldChanges. For the female groups, only log_2_FoldChange >0.5 or <−0.5 are shown. For the male groups, only log_2_FoldChange >1.0 or <−1.0 are shown. Color coding is based on functional enrichment analysis within STRING and is indicated for each treatment group. Abbreviations: GO-Gene Ontology, KEGG-Kyoto Encyclopedia of Genes and Genomes, ECM-extracellular matrix.

### Conservation of DEGS in Continuous Treatment Groups of Both Sexes

Additional analysis was done to compare the effects of the same treatment in different sexes (Figure 5). Venn diagrams (Figure 5a) illustrate that relatively few DEGs were common between the same treatment of different sexes, with the most overlap occurring between the high dose continuous groups. Interestingly, in the continuous dosing groups with low and high dose, four genes are conserved in both female and male cells indicating similar control of estrogen on the transcription of these genes. The proteins for which these genes encode function as facilitators of transport across cell membranes (*SLC7AA1 and ABCC1)*, protease that functions to cleave insulin-like growth factor binding proteins necessary for bone formation, inflammation, wound healing, and fertility (*PAPPA1 and 2)*, and transcription factor and ubiquitin binding (*UBXN7)*. The complete list of these overlapping DEGs can be found in Table S2. DAVID analysis (Figure 5b) revealed several groups enriched in these datasets compared to the human genome including extracellular exosomes in the low dose continuous treatments and extracellular space in the high dose continuous groups. STRING analysis (Figure 5c) illustrated that there were few protein-protein interactions between these genes. However, interactions controlling cell cycle processes (PAPPA/PCNA/TMSB4X) and TGFβR1/SMAD9, both important for meniscus cell proliferation and phenotype, were highlighted in the high dose continuous groups of both sexes.

**Figure 5:**
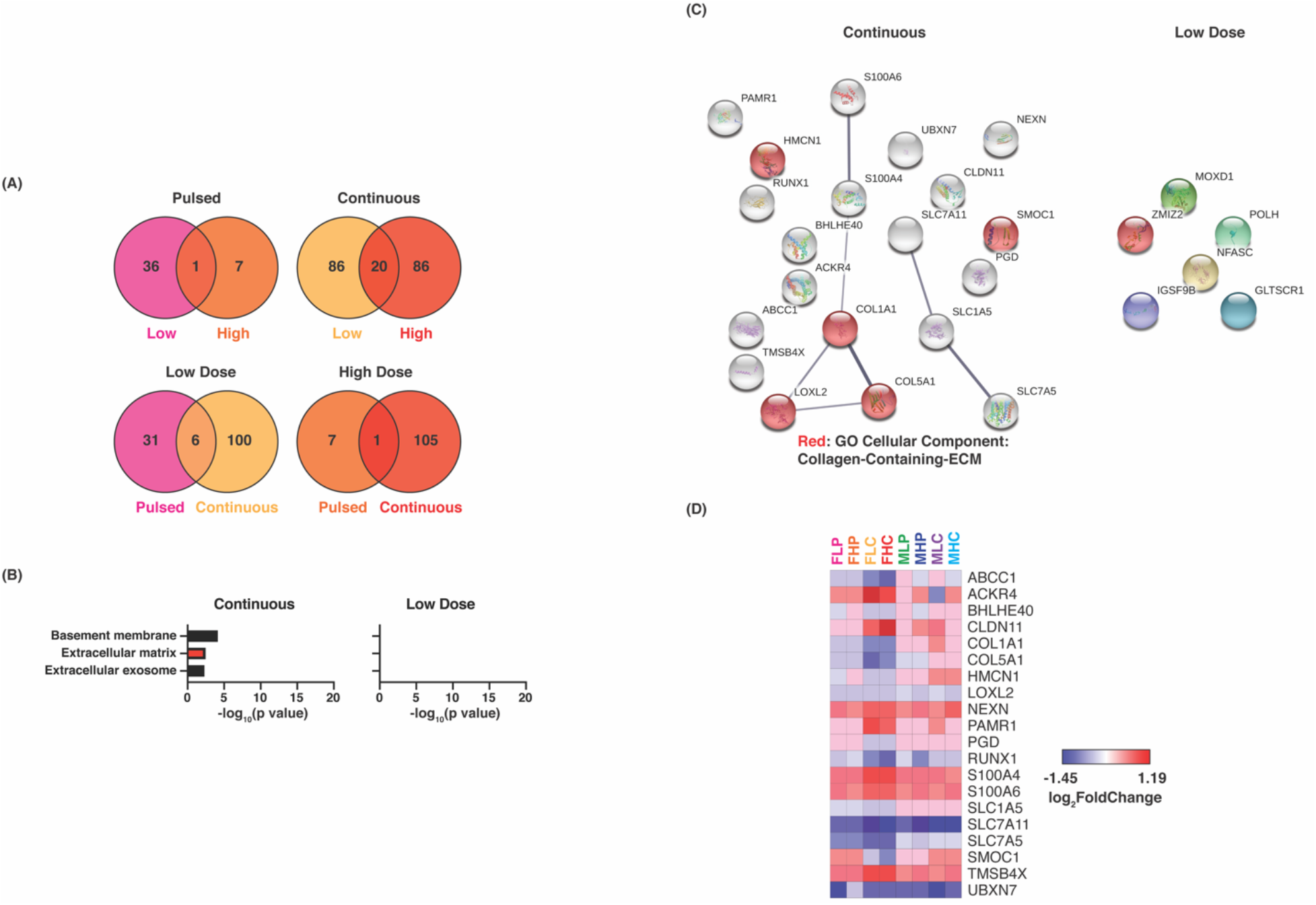
Analysis of the response of both sexes to the same treatment. (A) Venn diagrams (Venny 2.1) were generated to show which genes were altered in both sexes under the same treatment. For treatments with more than one common gene, functional enrichment analysis was performed using (B) DAVID and (C) STRING analysis to determine what types of genes were differentially expressed. Functional enrichment analysis shows the Gene Ontology Cellular Component groups with p values<0.05. STRING analysis shows known and predicted interactions between the proteins encoded by the genes in confidence views, where associations with stronger data support are thicker.

Overlapping DEGs as a function of treatment in the female groups are shown in Figure 6. Overall, continuous dosing exhibited the most conserved DEGs with estrogen concentration (Figure 6a). In looking at the pathway enrichment (Figure 6b) and protein-protein interactions (Figure 6c), genes involved in repair and homeostasis of the ECM are reported. All 20 DEGs of this group are shown with their expression in all other groups in the heat map in Figure 6d. The major genes regulated were all protein coding genes that are implicated in cell-cell signaling (*ACKR4, CLDN11)*, or cell differentiation (*PAMR1, S100A4, and S100A5*). A full list of these genes can be found in Table S3.

**Figure 6:**
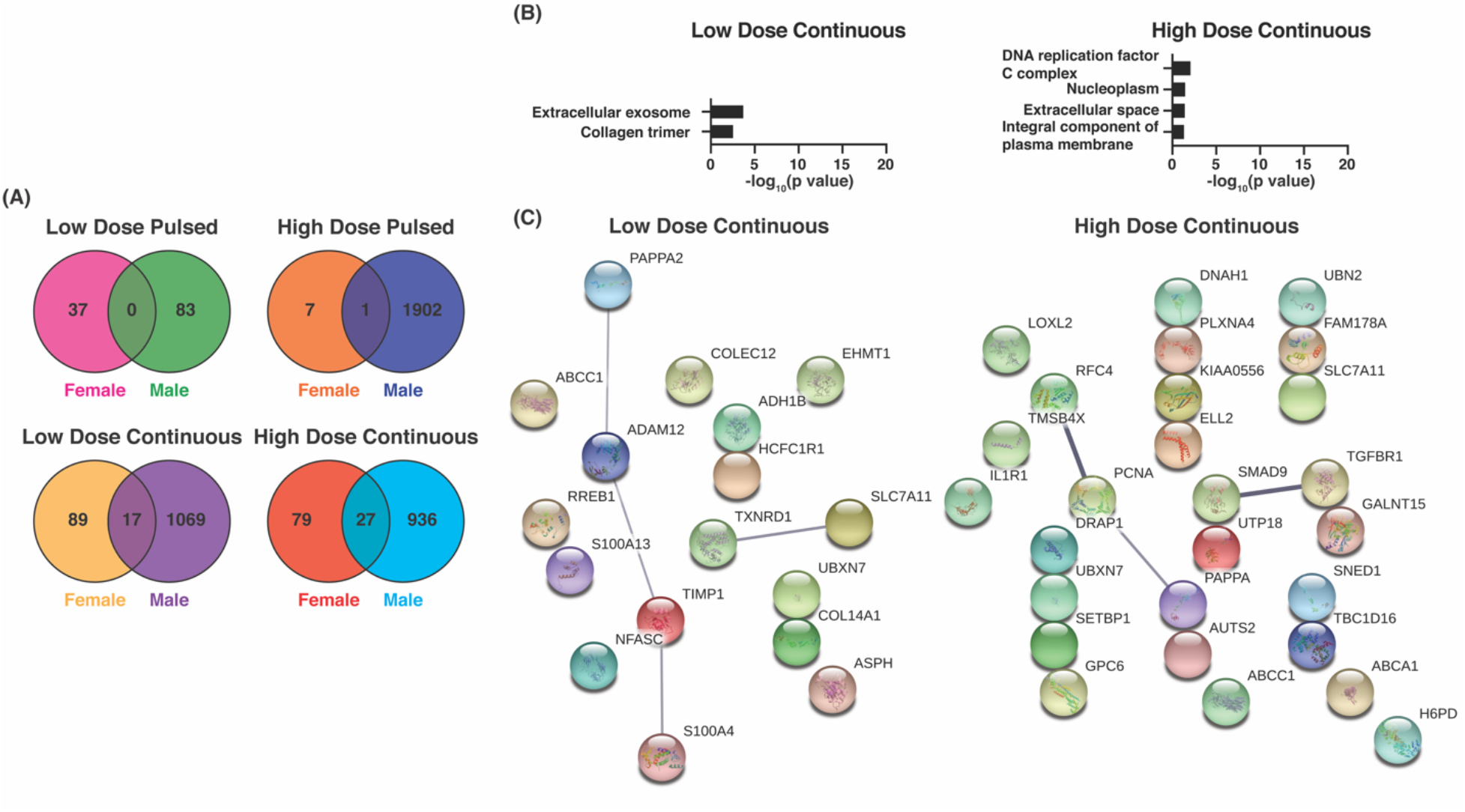
Analysis of DEGs that overlap between any two female treatment groups. (A) Venn diagrams (Venny 2.1) were generated to show which genes were altered in more than one treatment group. For groups with more than one common gene, functional enrichment analysis was performed using (B) DAVID and (C) STRING analysis to determine what types of genes were differentially expressed. (D) A heat map was generated of all 20 DEGs that were present in both continuous treatment groups. Functional enrichment analysis shows the Gene Ontology Cellular Component groups with p values<0.05. STRING analysis shows known and predicted interactions between the proteins encoded by the genes in confidence views, where associations with stronger data support are thicker. Heat map was generated using GenePattern. Genes are listed in alphabetical order. Values shown are log_2_FoldChange of each gene for each treatment group as compared to the sex-matched no E2 control.

### DEGS Related to the Extracellular Matrix from Estrogen Treatment

Functional enrichment analysis using DAVID^34^ showed that the types of genes enriched varied depending on donor sex, E2 dose, and dosing kinetics (Figure 7). Of interest for joint healing, genes of the ECM, highlighted in red, were enriched in all treatment groups except the female pulsed dosing groups. Genes related to extracellular exosomes and focal adhesion were enriched in all the male treatment groups plus the female low dose continuous group. Membrane, cytoplasm, and cytosol genes were enriched in all male treatment groups but none of the female treatment groups. Genes of the actin cytoskeleton were only enriched in both male low dose groups. Nucleus genes were enriched in all male groups except low dose pulsed. Genes of the basement membrane, extracellular region, extracellular space, collagen trimer, and endoplasmic reticulum lumen were enriched in both female continuous dosing groups but not in any of the male treatment groups.

**Figure 7:**
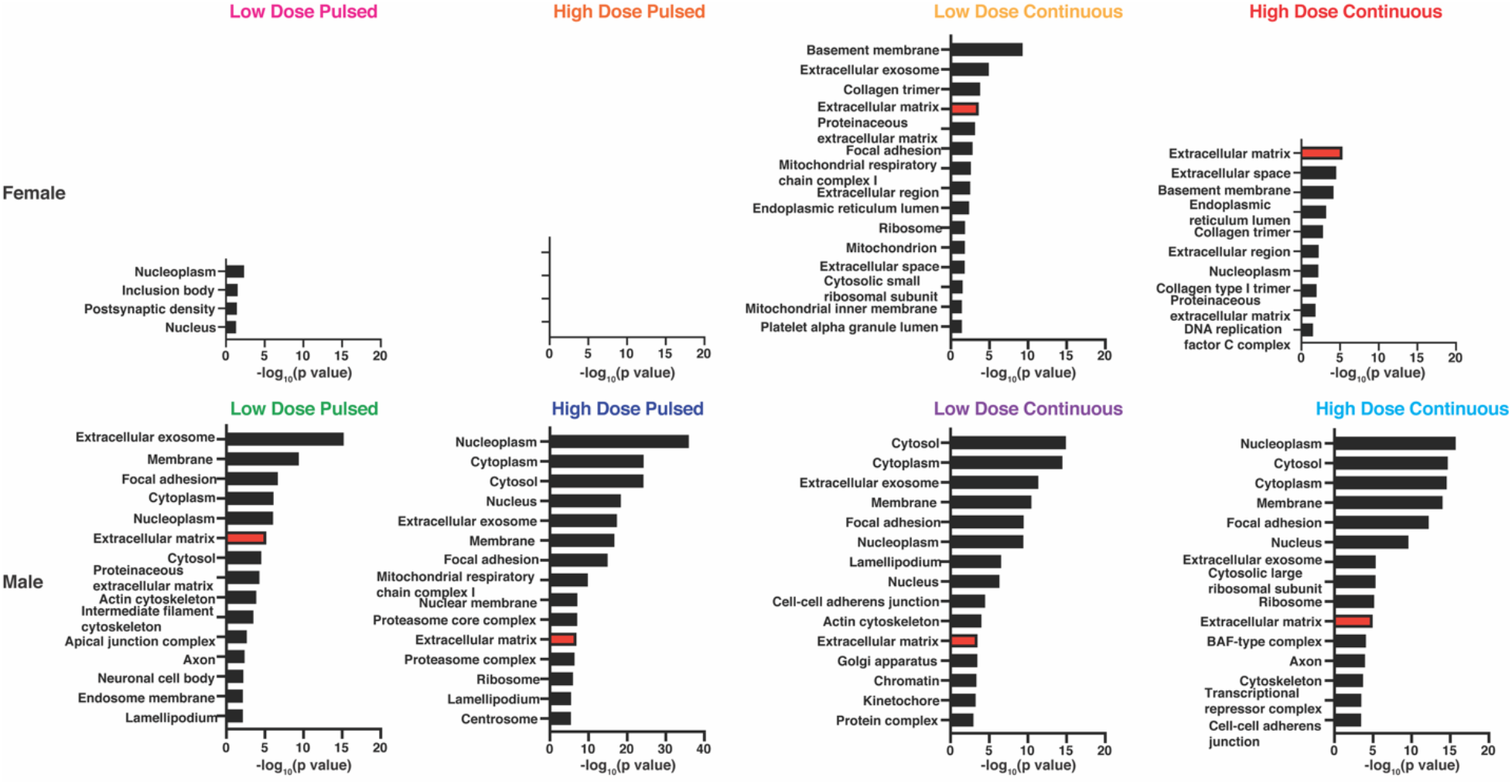
Functional enrichment analysis of DEGs using DAVID. Categories shown are from the Gene Ontology Cellular Component database. Where available, the top 15 groups by p value are shown with only p values < 0.05 considered.

The importance of estrogen in promoting new extracellular matrix for fibrocartilage tissue regeneration and homeostasis has been previously detailed.^36^ For this reason, the relative expression of genes within the ECM category from each treatment group are illustrated in a heat map in Figure 8. Separate heat maps for both sexes, doses, and dosing kinetics can be found in Figures S5-S7. The expression levels of many genes were either up- or down-regulated in all treatment groups. For example, ABI3BP, ADAMTS3, CANX, CDON, FBN1 and 2, LOXL2, LTBP2, TIMP2 and 3, and many collagens were down-regulated in all treatment groups and CALR, CFL1, CRIP2, LOXL1, LTBP3 and 4, and TNXB were up-regulated in all groups. Others such as ACTG1, COL14A1, COL7A1, HMCN1, mATN2, OGN, and PFKP were split by donor sex. No clear patterns were present based on low versus high dose, except for the ribosomal proteins in male treatment groups. Of particular interest to cartilage and fibrocartilage, COMP, POSTN, and TGFB1 and 3 were differentially regulated based on sex and estrogen dosing kinetics.

**Figure 8:**
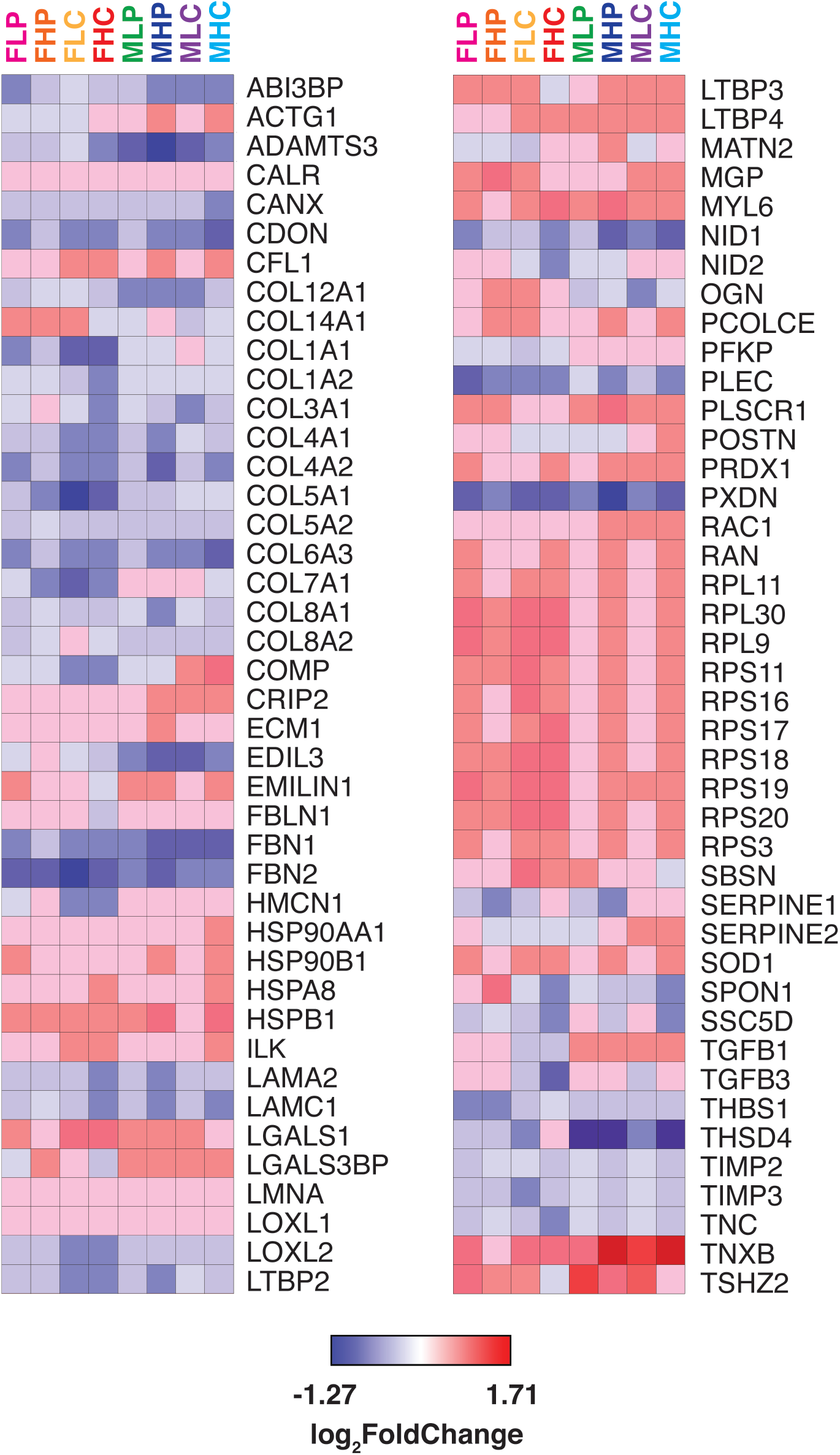
Heat map of extracellular matrix gene expression. Heat map was generated using GenePattern. Genes are listed in alphabetical order. Values shown are log_2_FoldChange of each gene for each treatment group as compared to the sex-matched no E2 control.

## Discussion

A number of microarray and sequencing studies have provided important insight on transcriptome differences in the meniscus including avascular meniscus that heals poorly,^37^ in meniscus from patients with and without osteoarthritis,^38^ the progenitor cell phenotype that may reduce osteoarthritis,^39^ and single-cell sequencing to define unique meniscal progenitor populations that are critical to maintain tissue health and reduce degeneration.^40^ Building off this body of work, this study focuses on transcriptome differences between male and female tissue that will inform sex differences in natural aging and inform the role of sex hormones in repair and homeostasis.

There is evidence that estrogen replacement therapy can reduce OA incidence and symptoms.^8–11^ ERT can also prevent and treat osteoporosis and osteoarthritis. Unfortunately, ERT is not without risk and can have negative off-target effects. Oral use of ERT increases the risk of venous thromboembolism in post-menopausal women.^41^ Combination use of estrogen plus progestin, and to a lesser extent estrogen alone, is linked with an increased risk of breast cancer in some studies.^42; 43^ The results presented here contribute to the pool of evidence supporting the importance of estrogen dose and kinetics for achieving the desired results in different patient populations. Further studies could lead to knowledge about estrogen dose and kinetics that are needed to develop safe and effective treatments for knee injury and mitigate the need for systemic dosing at high concentrations needed to hit therapeutic dose that lead to deadly, off-target effects.

While more DEGs were reported in all of the male treatment groups, a hypothesized effect due to the relative naivety of the male cells to high estrogen doses, an important finding is the interplay between concentration and kinetics in both groups. In most cases, there were more DEGs seen in treatment groups with continuous dosing than with pulsed dosing. This corresponds with the findings of Simoncini *et. al.* that continuous E2 dosing of human umbilical vein endothelial cells (HUVECs) leads to heavier E2 signaling through the traditional genomic pathways than pulsed dosing.^16^ In contrast, Cavailles *et. al.* found no difference in the effects of pulsed compared to continuous estrogen dosing on the expression levels of 3 estrogen-responsive genes.^20^ This inconsistency can possibly be explained by the different scopes of the studies and different concentrations of E2 used. Another study found more DEGs in HUVECs treated with pulsed estrogen than those treated with continuous estrogen,^17^ in direct contradiction to the current study. A possible explanation for this is the differential response to estrogen dosing kinetics based on tissue type seen by Otto *et. al.^18^* Interestingly, with continuous dosing, concentration did not play a major role in total DEGs whereas concentration did play a significant role with pulsed dosing. This evidence may indicate the ability to achieve similar transcriptional effects using continuous dosing at lower concentrations which is important to reduce the negative, off-target effects to other tissues in ERT. Also, in the female groups, pulsed dosing resulted in lower total DEGs whereas the total DEGs was a function of concentration in the male groups. Overall, it is likely that continuous dosing is required for the robust transcriptional changes necessary for remodeling and homeostasis to maintain meniscal health and reduce OA.

Functional enrichment analysis using DAVID and STRING revealed that the types of genes enriched varied by donor sex, E2 dose, and dosing kinetics. Genes related to the extracellular matrix were enriched in all groups except the female pulsed groups. It is also of interest that the DEGs common to both male and female cells are related to the extracellular matrix. While all of the differentially expressed collagens and tissue inhibitors of metalloproteinases (TIMPs) are downregulated within all treatment groups, growth factors such as in the TGFβ family are upregulated in all male groups and in the female pulsed groups. This is information that must be pursued in future studies as TGFβ3 is a major growth factor in meniscus remodeling and repair^44–46^ and activation of this protein enhances differentiation ability of meniscus progenitors to chondrogenic-like avascular tissue.^39; 47^ Additionally, TNXB, LTBP3, and LTBP4, genes that have each been shown to play a role in the activation, availability, and/or signaling of TGFβ^48; 49^, are all upregulated in all groups. LTBP2, another gene in the LTBP family, is down-regulated, but it is also the only one in the family that does not bind to latent TGFβ.^49^ TNXB is also important for the correct assembly of other matrix molecules and wound healing.^49^ Further, this study is only assessing transcriptional changes after 72 hours and thus is likely not capturing all of the transcriptional changes occurring with E2 treatment.

There are limitations to the current study that should be considered. This study was performed using 2D culture methods, which are known to promote a more fibroblast-like phenotype of the meniscal fibrochondrocytes^37; 50^ and thus will indicate more of the fibroblast-like cell response to estrogen. Additionally, this study was performed using cells isolated from only 2 donors, both 47-years old. It is likely that these cells had relatively limited remodeling capability due to the age of the donors. Furthermore, the 2 donors were of different races (Black and Caucasian), which could have been a complicating factor. Future studies on 3D cultures using tissue from more donors of different ages and more concentrations of estrogen could be very informative to elucidate more conclusive effects of estrogen on the meniscus and for the development of treatments for knee injury.

## Conclusions

Overall, the estrogen dosing kinetics played a more significant role on total DEG compared to estrogen concentration in both sexes. There were only four DEGs conserved across sexes in both low and high concentrations in continuous dosing illustrating that meniscus cell transcriptional response to estrogen signaling is significantly different between male and females. Future work will expand on this initial data set to probe the effect of specific pathways, such as TGF-β3 known to promote meniscus healing, to provide clear insight into the role estrogen plays on meniscus regeneration and homeostasis. Overall, this initial study lays the groundwork for future avenues to pursue the effect of estrogen delivery on regenerative pathways to inform the design and implementation of ERT therapies to promote meniscal health and reduce the onset of osteoarthritis.

## Acknowledgments

The authors acknowledge the use of tissues procured by the National Disease Research Interchange (NDRI) with support from NIH grant U42OD11158. The authors want to thank Brittney Smith and Jennifer Hackett at the KU Genome Sequencing Center (GSC) for completing the sequencing (NIH NIGMS P20GM103638) and Boryana Koseva and Stuart Macdonald of the K-INBRE Bioinformatics Core for providing computational support (NIH P20 GM103418). Further, the authors are appreciative of the insight and analysis feedback from Dr. Emily Farrow, director of laboratory operations at the Genome Center at Children’s Mercy Hospital. This work was supported by startup funds from the University of Kansas (JR) and NIH NIGMS P20GM103638 (KK, JGG, and JLR). This manuscript appears in the pre-print server bioRxiv (2020.04.27.064451, https://www.biorxiv.org/content/10.1101/2020.04.27.064451v1).

