## Supplementary Information for "Transcriptome Sequencing Reveals Sex Differences in Human Meniscal Cell Response to Estrogen Based on Dosing Kinetics"

| | | | | | |
|---|---|---|---|---|---|
| ABCC1 | CDC42BPA | ERBIN | LIFR | PIK3R1 | STAG2 |
| ABI3BP | CEMIP | ERC1 | LMAN1 | PIKFYVE | SVEP1 |
| ABLIM1 | CEP350 | EZH1 | LNPEP | PRKAR2A | TAOK1 |
| ACAP2 | CFAP97 | FAM91A1 | LRRC58 | PRPF8 | TAP1 |
| ADCY9 | CHD2 | FAT4 | LTN1 | PRR14L | TBCD |
| AFAP1 | CLASP1 | FBN1 | LYST | PRRC2B | TBX18 |
| AFF4 | CLIC3 | FBN2 | MAN1A1 | PSMB1 | TCAF1 |
| AHNAK | CLIC4 | FNIP1 | MAN2A1 | PSMD4 | TEAD1 |
| AHR | CLOCK | FRY | MDH2 | PTBP3 | TFRC |
| AKAP11 | COL6A3 | FXYD5 | MDM2 | PTPRG | TGFB1 |
| ALCAM | COX5B | GADD45GIP1 | MDN1 | PUM2 | TGFBR1 |
| ALDH1L2 | CPD | GNAQ | MED13 | PXDN | TIMP1 |
| ANKRD52 | CPSF1 | GPATCH8 | MED13L | QKI | TJP1 |
| ANO6 | CREB3L2 | GPC6 | MGAT4B | RAB3GAP2 | TMTC3 |
| AP4E1 | CREBRF | GSTO1 | MIB1 | RALGAPB | TNKS2 |
| APPL1 | CRIM1 | GTF2A1 | MRPS24 | RAPH1 | TNRC6A |
| ARFGEF2 | CRIP1 | H6PD | MYL6 | RERE | TNRC6C |
| ARHGAP29 | CRTC3 | HCFC1R1 | MYSM1 | RIC1 | TNXB |
| ARHGAP35 | CUTA | HECA | N4BP2 | RIF1 | TPCN1 |
| ARHGEF12 | DAB2 | HEG1 | NBEA | ROBO1 | TRPS1 |
| ARL6IP4 | DDR2 | HERC2 | NF1 | ROCK2 | UHMK1 |
| ARPC1B | DDX17 | HIPK1 | NFE2L1 | SASH1 | VARS |
| ASAP1 | DDX6 | HIPK3 | NFIB | SCN8A | VCPIP1 |
| ASH1L | DICER1 | HIVEP2 | NID1 | SEC14L1 | VPS13A |
| ASXL2 | DIP2B | HSPB1 | NOTCH2 | SECISBP2L | VPS13C |
| ATIC | DIXDC1 | IFITM2 | NR1D2 | SEL1L | WNK1 |
| ATM | DMXL1 | IFITM3 | NR3C1 | SEMA3C | XRN1 |
| ATP11B | DNAJC3 | IGFBP5 | NUFIP2 | SETD7 | ZBED6 |
| ATP9A | DNAJC7 | ITGA11 | OSBPL8 | SLC12A2 | ZBTB41 |
| ATRN | DOCK1 | ITGAV | OSMR | SLC38A1 | ZDHHC20 |
| BAHCC1 | DPYSL2 | JADE1 | OTUD4 | SLC38A2 | ZFP91 |
| BICRAL | DRAP1 | JMY | PAPPA2 | SLC39A10 | ZNF407 |
| BRCA2 | EDIL3 | KDM2A | PBRM1 | SLC39A14 | ZNF609 |
| CAMSAP2 | EEA1 | KIAA0232 | PDGFRA | SLC7A11 | ZNF770 |
| CANX | EGFR | KIF1B | PDLIM7 | SMAD4 | ZNF827 |
| CBLL1 | EMILIN1 | KIRREL1 | PEAK1 | SMG7 | |
| CCDC186 | ENAH | KMT2C | PHC3 | SNTB2 | |
| CD109 | EP300 | KNL1 | PIEZO2 | SOS1 | |
| CD81 | EPAS1 | LATS1 | PIK3C2A | SPRED2 | |

**Table S1:** List of DEGs common to all male treatment groups. Genes are listed alphabetically.

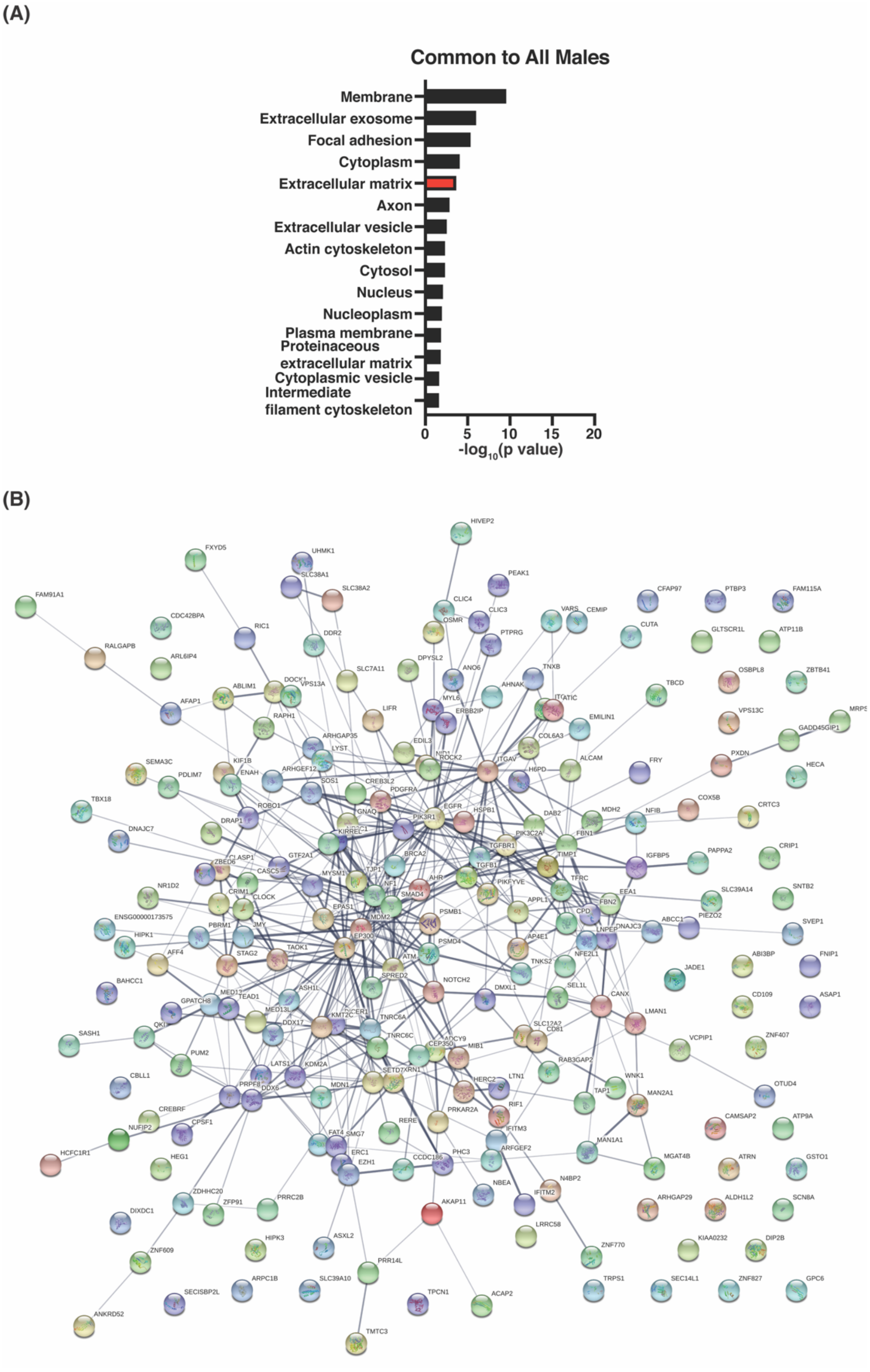


**Figure S1:** Analysis of DEGs common to all male treatment groups. Functional enrichment analysis performed using DAVID (A) and STRING (B) reveal the types of differentially expressed genes and their interactions. Functional enrichment analysis shows the Gene Ontology Cellular Component groups with p values<0.05. STRING analysis shows known and predicted interactions between the proteins encoded by the genes in confidence view, where associations with stronger data support are thicker.

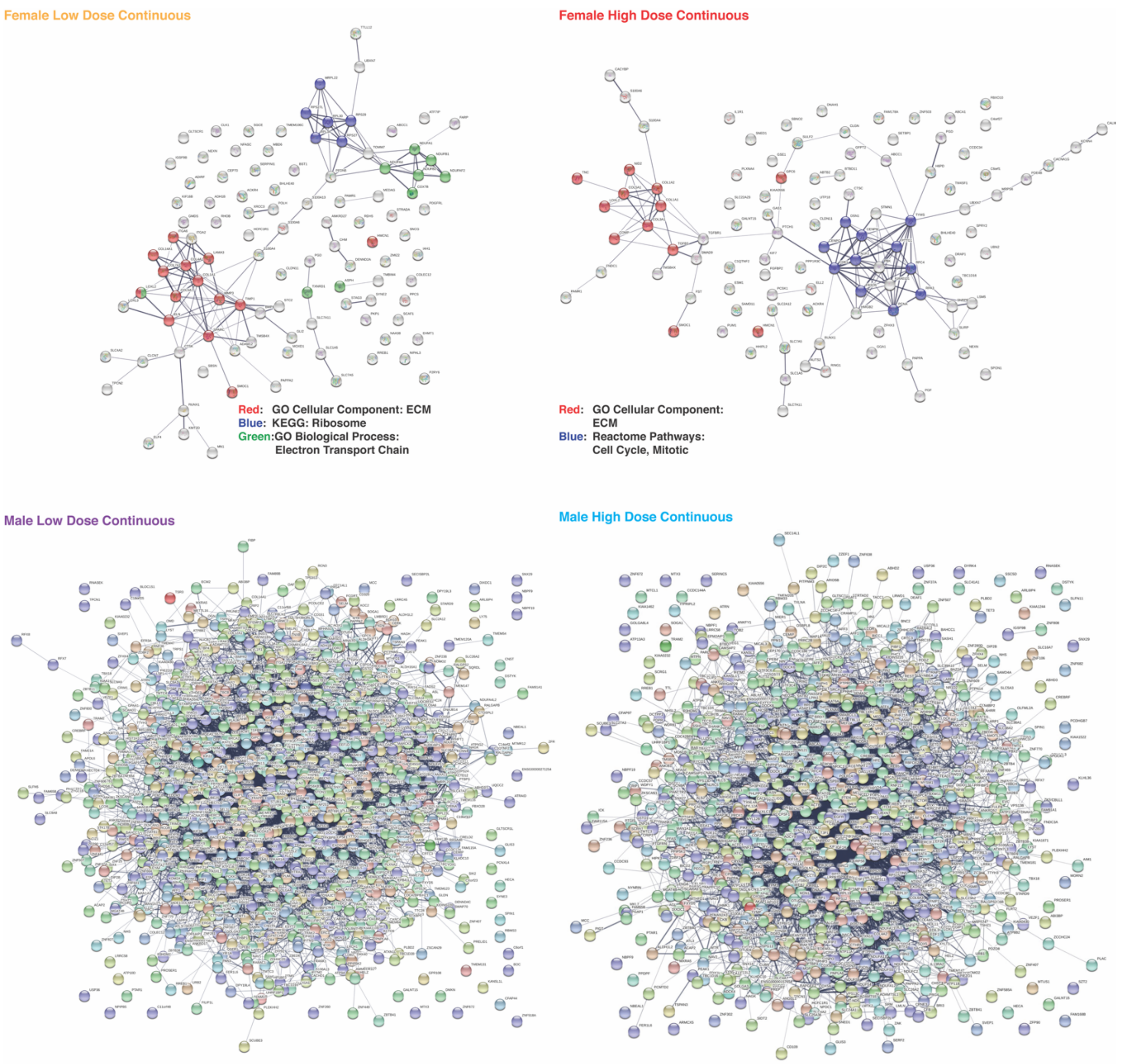


**Figure S2:** STRING analysis of DEGs from all 4 continuous dosing treatment groups. Images are shown as confidence views, where associations with stronger data support are thicker. Only DEGs with padj<0.05 are shown. Color coding in the female groups is based on functional enrichment analysis within STRING and is indicated for each treatment group. Abbreviations: GO-Gene Ontology, KEGG-Kyoto Encyclopedia of Genes and Genomes, ECM-extracellular matrix

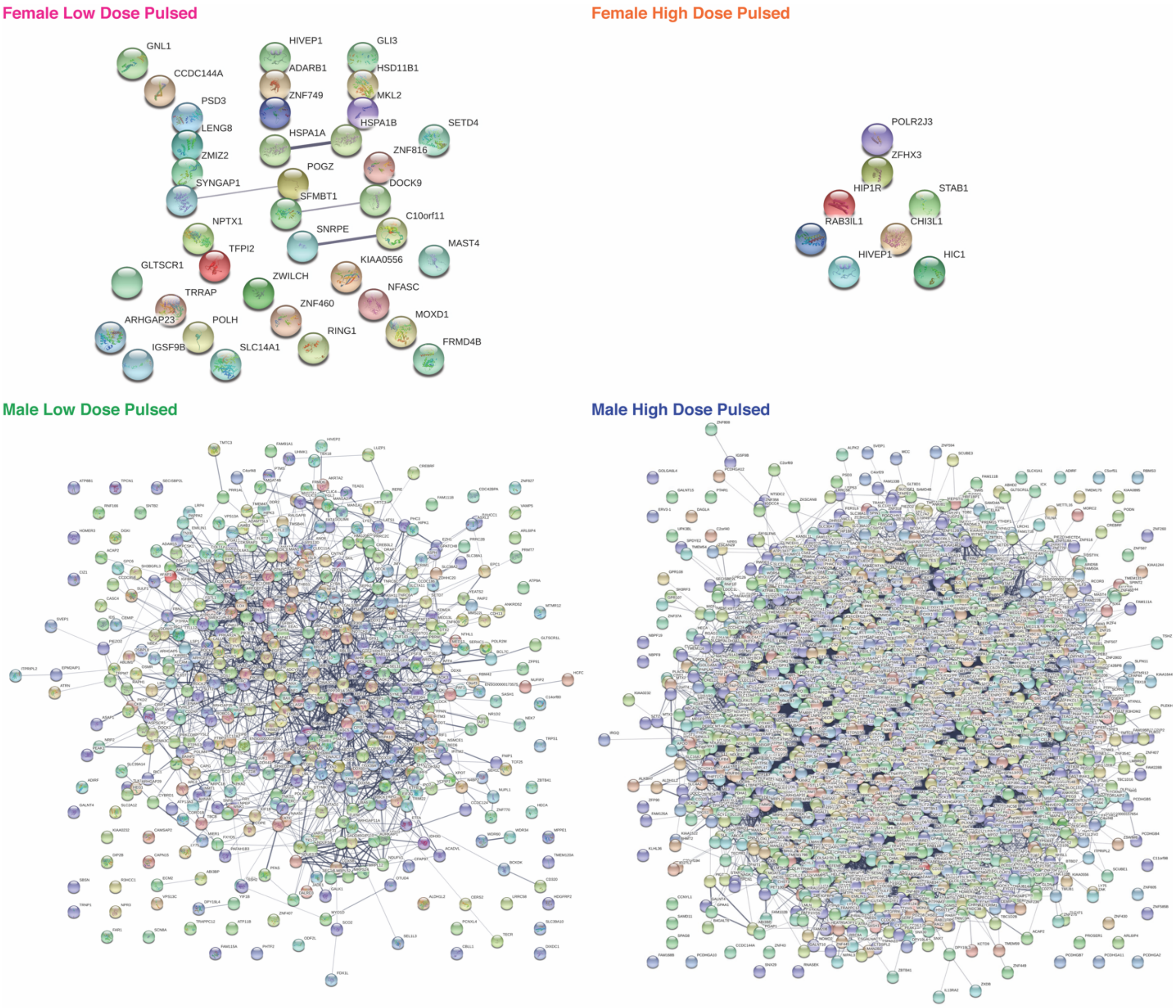


**Figure S3:** STRING analysis of DEGs from all 4 pulsed dosing treatment groups. Images are shown as confidence views, where associations with stronger data support are thicker. Only DEGs with padj<0.05 are shown.

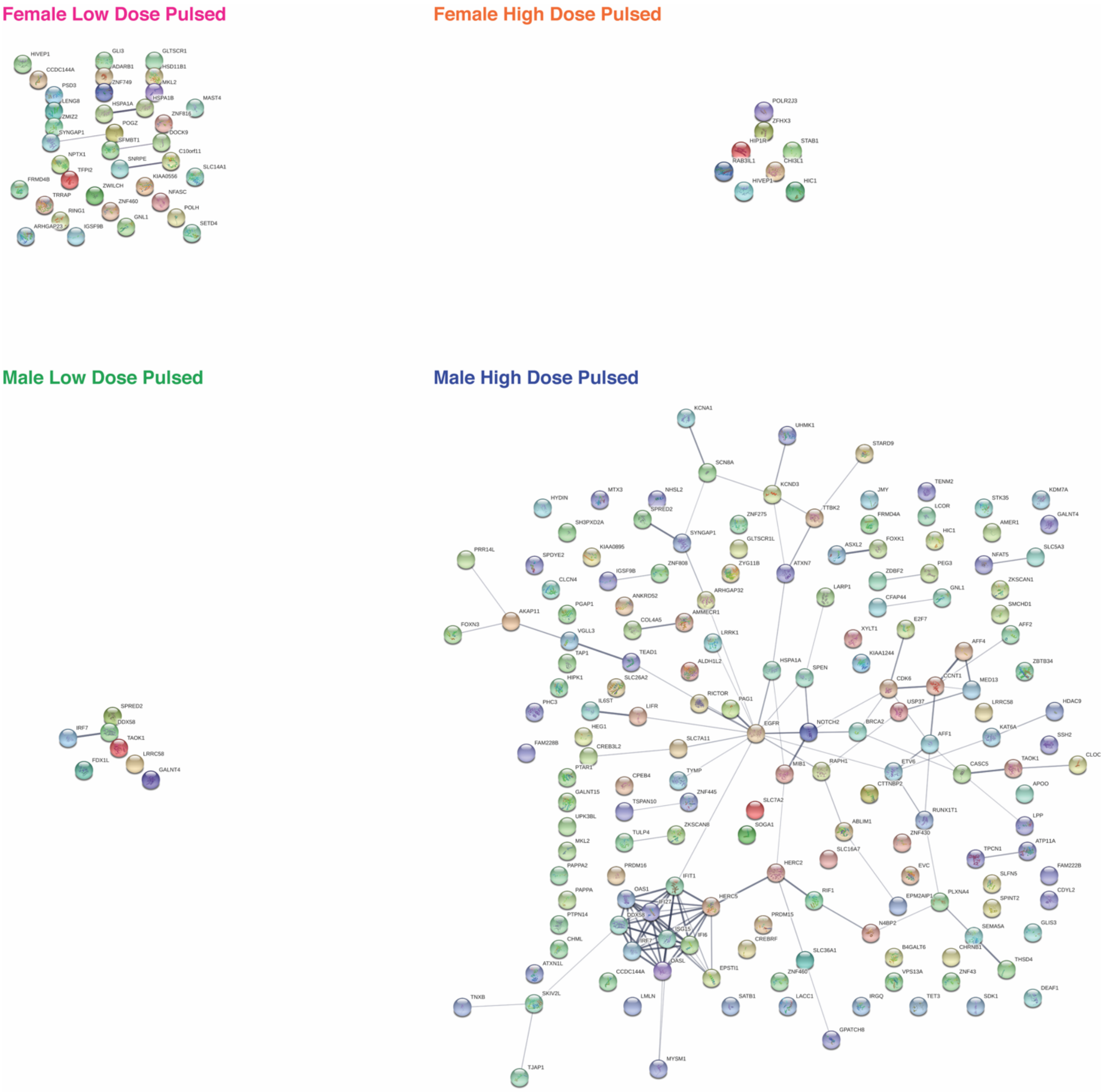


**Figure S4:** STRING analysis of DEGs from all 4 pulsed dosing groups. Images are shown as confidence views, where associations with stronger data support are thicker. Only DEGs with padj<0.05 are shown. Additional constraints based on $log_2$FoldChange were used to match the limits applied in Fig 4 of the main body of the manuscript. For the female groups, only $log_2$FoldChange >0.5 or <-0.5 are shown. For the male groups, only $log_2$FoldChange >1.0 or <-1.0 are shown.

| **LDP** | **HDP** | **LDC** | **HDC** |
|---|---|---|---|
| | HIC1 | SLC7A11 | PAPPA |
| | | TXNRD1 | SNED1 |
| | | ASPH | RFC4 |
| | | COLEC12 | SETBP1 |
| | | ADAM12 | ABCA1 |
| | | NFASC | PLXNA4 |
| | | RREB1 | KIAA0556 |
| | | S100A4 | ABCC1 |
| | | EHMT1 | SLC7A11 |
| | | ABCC1 | UBN2 |
| | | ADH1B | SLF2 |
| | | S100A13 | AUTS2 |
| | | COL14A1 | DRAP1 |
| | | PAPPA2 | TGFBR1 |
| | | UBXN7 | TMSB4X |
| | | TIMP1 | GALNT15 |
| | | HCFC1R1 | TBC1D16 |
| | | | DNAH1 |
| | | | H6PD |
| | | | IL1R1 |
| | | | UBXN7 |
| | | | PCNA |
| | | | SMAD9 |
| | | | UTP18 |
| | | | LOXL2 |
| | | | GPC6 |
| | | | ELL2 |

**Table S2:** List of DEGs seen in both sexes by treatment.

| **Pulsed** | **Continuous** | **Low Dose** | **High Dose** |
|---|---|---|---|
| HIVEP1 | SMOC1 | IGSF9B | ZFHX3 |
| | COL5A1 | POLH | |
| | PAMR1 | BICRA | |
| | CLDN11 | ZMIZ2 | |
| | NEXN | NFASC | |
| | COL1A1 | MOXD1 | |
| | RUNX1 | | |
| | ACKR4 | | |
| | ABCC1 | | |
| | SLC7A11 | | |
| | PGD | | |
| | TMSB4X | | |
| | SLC7A5 | | |
| | SLC1A5 | | |
| | BHLHE40 | | |
| | HMCN1 | | |
| | UBXN7 | | |
| | LOXL2 | | |
| | S100A4 | | |
| | S100A6 | | |

**Table S3:** List of DEGs seen in two female treatment groups.

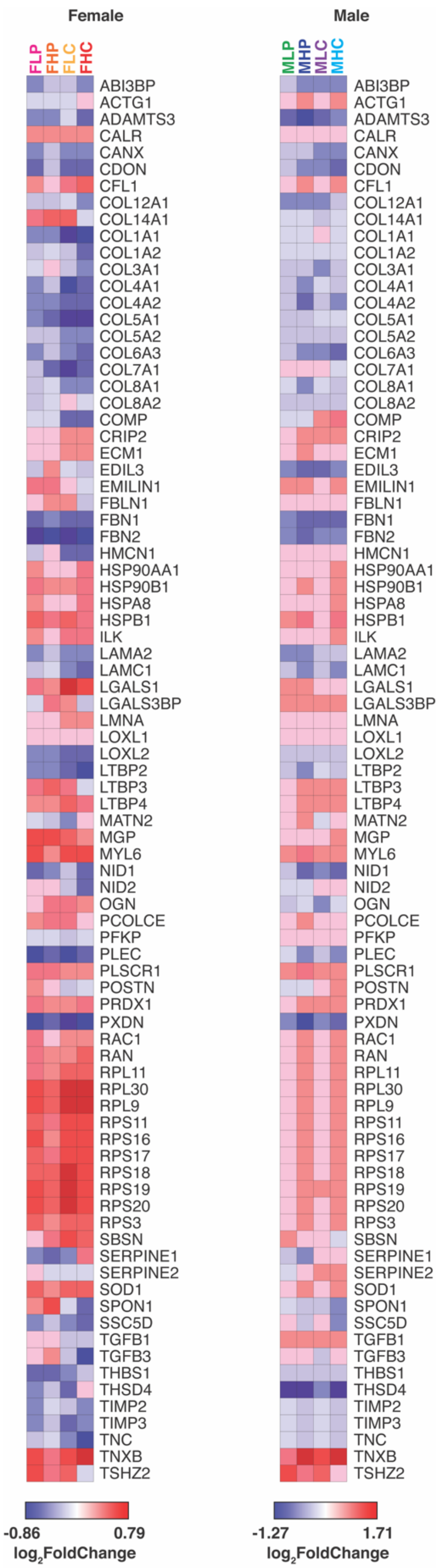


**Figure S5:** Heat maps of extracellular matrix gene expression split by sex. Heat maps were generated using GenePattern. Genes are listed in alphabetical order. Values shown are $\log_2$FoldChange of each gene for each treatment group as compared to the sex-matched no E2 control.

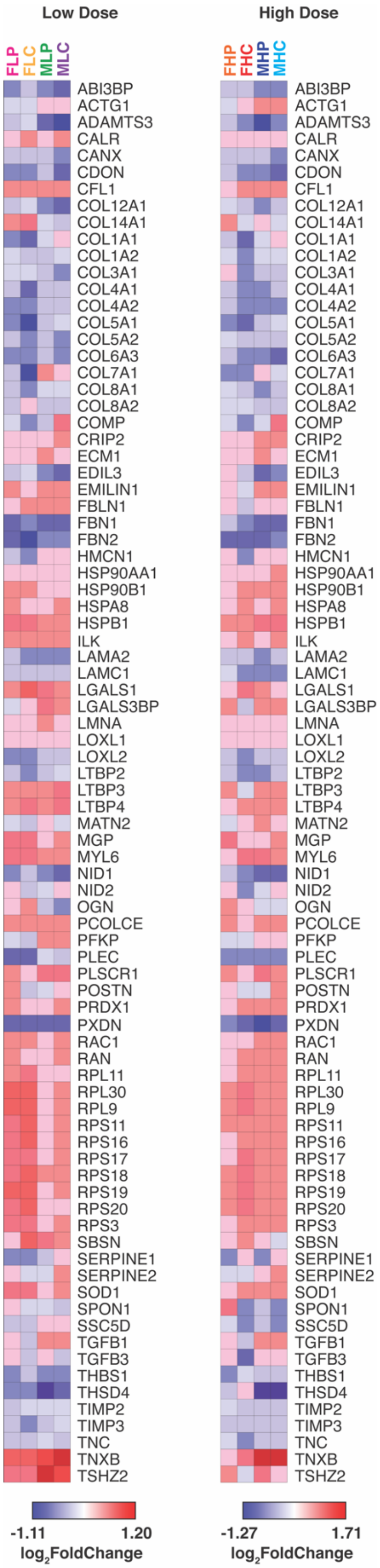


**Figure S6:** Heat maps of extracellular matrix gene expression split by estrogen dose. Heat maps were generated using GenePattern. Genes are listed in alphabetical order. Values shown are $\log_2$FoldChange of each gene for each treatment group as compared to the sex-matched no E2 control.

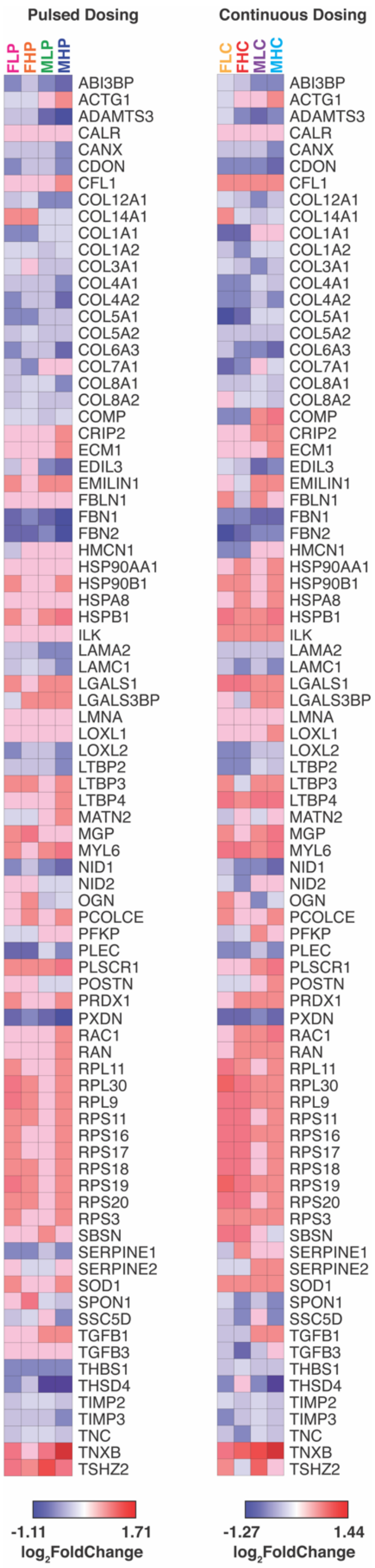


**Figure S7:** Heat maps of extracellular matrix gene expression split by dosing kinetics. Heat maps were generated using GenePattern. Genes are listed in alphabetical order. Values shown are $\log_2$FoldChange of each gene for each treatment group as compared to the sex-matched no E2 control.
